# Comparison of Machine Learning Surrogate Models for Prediction of Single-Fiber Activation in Deep Brain Stimulation

**DOI:** 10.64898/2026.05.12.724686

**Authors:** Jorge Alberto, Benjamin Norbom, Justin Golabek, Joshua K. Wong, Matthew Schiefer, Erin E. Patrick

## Abstract

Machine-learning surrogate models are positioned to help optimize deep brain stimulation (DBS) usage by predicting neural activation in response to electrical stimulation, while minimizing tradeoffs between computational expense and accuracy. Previous work has developed high accuracy artificial neural network (ANN) and convolutional neural network (CNN) surrogate models that predict activation of individual, myelinated axons, to extracellular electrical stimulation for subsets of DBS programming configurations. Moreover, more traditional machine learning methods including extreme gradient boosting (XGBoost) have been used effectively for peripheral-nerve single-fiber activation predictions. We build upon the previous work and compare ANN, CNN and XGBoost methods to a much expanded set of electrode programming configurations including: monopolar, bipolar, tripolar, quadrupolar, multiple monopolar, and multiple cases of directional leads. Training used datasets generated from a finite-element model of an implanted DBS lead together with multi-compartment cable models of synthetically generated axons. We evaluated the machine learning predictors using white matter pathways derived from group-averaged connectome data within a patient-specific tissue conductivity field, comparing both predicted stimulus activation thresholds and pathway recruitment across a clinically relevant range of stimulus amplitudes and pulse widths. Our ANN and CNN models successfully predicted neural fiber activation for almost all electrode configurations with low error, expanding the scope of our previous predictor model. Results also showed key limitations of XGBoost models and superior performance of CNNs for more complex electrostatic fields of the directional leads.

## 1. Introduction

Deep Brain Stimulation (DBS) is a surgical procedure that delivers electrical stimulation to deep brain regions, resulting in therapeutic benefits for a range of neurological and psychiatric disorders [1]. While highly successful at over 300,000 total implants [2], DBS still heavily relies on a manual process for parameter optimization, where iterative adjustments to electrode configuration, frequency, amplitude, and pulse width are made by physicians [3]. Further slowing down this manual process, recent DBS innovations such as directional leads, anodic programming, and fractional current [4] have significantly increased the number of potential parameter combinations through combinatorial expansion, making the programming process further complex and time-consuming [5].

Computational methods can reduce this burden. In fact, Boston Scientific (BSC) has a commercially available system that allows clinicians to designate a volume of tissue activated (VTA), after which the programming software determines the required electrode configuration (i.e., which electrode/s on the DBS lead is/are active and the level of current or voltage stimulation) [5]. In addition to post-surgical programming, presurgical targeting [6] and retrospective studies that enable population-based statistics for disease-specific treatments (e.g., [7–13]) employ computational models for prediction of DBS activation.

While computational models hold promise for addressing challenges to presurgical targeting and postsurgical programming, which need near instant predictive results when used clinically, they face a fundamental trade off: models with high biophysical accuracy often come with significant computational complexity. In particular, models that possess finite element methods (FEM) to compute voltage fields in conjunction with simulation of multicompartment axon models and realistic pathways yield the highest biophysical accuracy but are extremely time consuming [14].

Thus, there has been a surge in single-fiber machine-learning surrogate models trained for DBS and peripheral nerve activation prediction. First, Golabek et al. addressed this computational expense by introducing an artificial neural network (ANN)-based surrogate model for activation of 5.7 mm diameter myelinated axons by DBS [15]. Their model reduced runtime by four to five orders of magnitude while preserving high predictive accuracy for realistic fiber pathways in anisotropic conductivity brain tissue models, demonstrating that surrogate modelling could bridge the gap between biophysical fidelity and computational efficiency. Despite these advances, the ANN was limited to prediction on monopolar electrode configurations and could not predict activation for directed leads and multipolar configurations, which are becoming increasingly common in clinical practice. Wang et al. subsequently extended the surrogate modelling framework to account for bipolar and segmented (directional) leads [16]. They also introduced convolutional neural networks (CNNs) by replacing some of the fully-connected layers of the ANN with 1D-convolutional layers. Both ANN and CNN models performed well but were only tested on straight axons and isotropic brain conductivity. This group’s follow-on work [17] presented another CNN-based model that could account for non-straight, realistic fiber pathways of variable diameter, although the results only showed examples in isotropic brain tissue models. Additionally, Toni et al. have innovated surrogate models for the peripheral nervous system [18]. They shifted away from purely exploring neural networks and also explored more traditional machine learning methods including eXtreme Gradient Boosting (XGBoost), support vector machine, and random forest. Their analyses focused on peripheral nerve stimulation of modelled, straight axonal fibers and found that XGBoost slightly outperformed an artificial neural network (“multilayer perceptron”) in terms of both accuracy and prediction time [18].

In this study, we extend the surrogate modelling approach for DBS introduced by Golabek et al. and Wang et al. by expanding ANN- and CNN-based predictors to model nerve fiber activation for not only directional leads, but also bipolar, multipolar electrode configurations and asymmetric, or fractional, stimulus amplitudes. We further enhance clinical applicability by explicitly evaluating out-of-distribution performance through testing unseen electrode configurations and realistic fiber tracts within an anisotropic conductivity field. In addition, we build on the work of Toni et al. by evaluating classical machine learning methods, particularly XGBoost, to determine whether their success in peripheral nerve stimulation translates to DBS, comparing XGBoost to ANN-based models in terms of accuracy, generalization, and computational efficiency.

## 2. Methods

The XGBoost workflow consisting of training and testing is presented first in section 2.2 and was used primarily to compare XGBoost machine-learning methods with our prior ANN model [15] to determine if tree-based methods could give as good accuracy for DBS models as those seen in Toni et al. [18] for the peripheral nervous system. The ANN and CNN methods are presented together in section 2.3, because both were designed to advance generalizability of our prior ANN model and are later directly compared against each other. Despite this separation, both models shared core components: the same volume conductor and axon model generation pipeline and a common evaluation procedure presented next.

### 2.1 Volume Conductor and Axon Models

The volume conductor and axon models in this study were generated following the same procedure as described by Golabek et al. [15], with the exception of using an updated COMSOL version (6.2) to create finite-element models (FEMs) of not only an implanted Medtronic 3387 DBS lead (Medtronic, Minneapolis, MN), but also the Boston Scientific Vercise Cartesia Directional DBS lead DB-2202-45 (Boston Scientific, Marlborough, MA). In summary, we used COMSOL 6.2 with the AC/DC module to solve Laplace’s equation for a given DBS lead in a 60 × 60 × 60 mm cube of bulk tissue, surrounded by 0.5 mm encapsulation tissue (σ = 0.07 S m^−1^) [19]. A quartic discretization was used in the FEM to enable calculation of accurate second spatial derivatives. The FEM had a minimum element size of 0.24 mm; mesh refinement ceased when voltages changed <1 % and the convergence criterion was 0.001. Training data were generated with homogeneous, isotropic tissue (σ = 0.2 S m^−1^) [20,21]; test data used a patient-specific anisotropic conductivity tensor from diffusion tensor imaging data [22], interpolated from 1 mm^3^ voxels. FEM potentials were exported on a 0.25 mm 3-D grid and interpolated with SciPy’s RegularGridInterpolator. Assuming quasi-static, purely resistive tissue [23], the FEM solution was linearly scaled for different stimulation amplitudes.

Axon models were multi-compartment double-cable 5.7 µm-diameter fibers built with the McIntyre–Richardson–Grill (MRG) model [24] in NEURON [25], capped with passive nodes to avoid phantom activations. Extracellular potentials at axonal compartments were obtained from the FEM solution, multiplied by a pulse train (monophasic cathodic), and applied via NEURON’s play method with a 5 µs timestep. Threshold voltages (Vth) – the minimum stimulus needed for neural activation– were determined to ±1 mV precision via binary search [20,26,27].

Synthetic fiber trajectories with a high degree of tortuosity were generated for the training and validation phases using a random walk algorithm that iteratively constructed pathways by connecting line segments with random directionality (up to a 30º angle) and smoothed with cubic splines following [15]. Realistic, non-straight trajectories for the testing phase were produced by extracting three white-matter pathways from the Human Connectome Project (1065 subjects) [28–31] for ground-truth testing. Graphical representations of the fiber tracts are shown in Figure 1. Fibers intersecting the lead or scar tissue were excluded.

**Figure 1.**
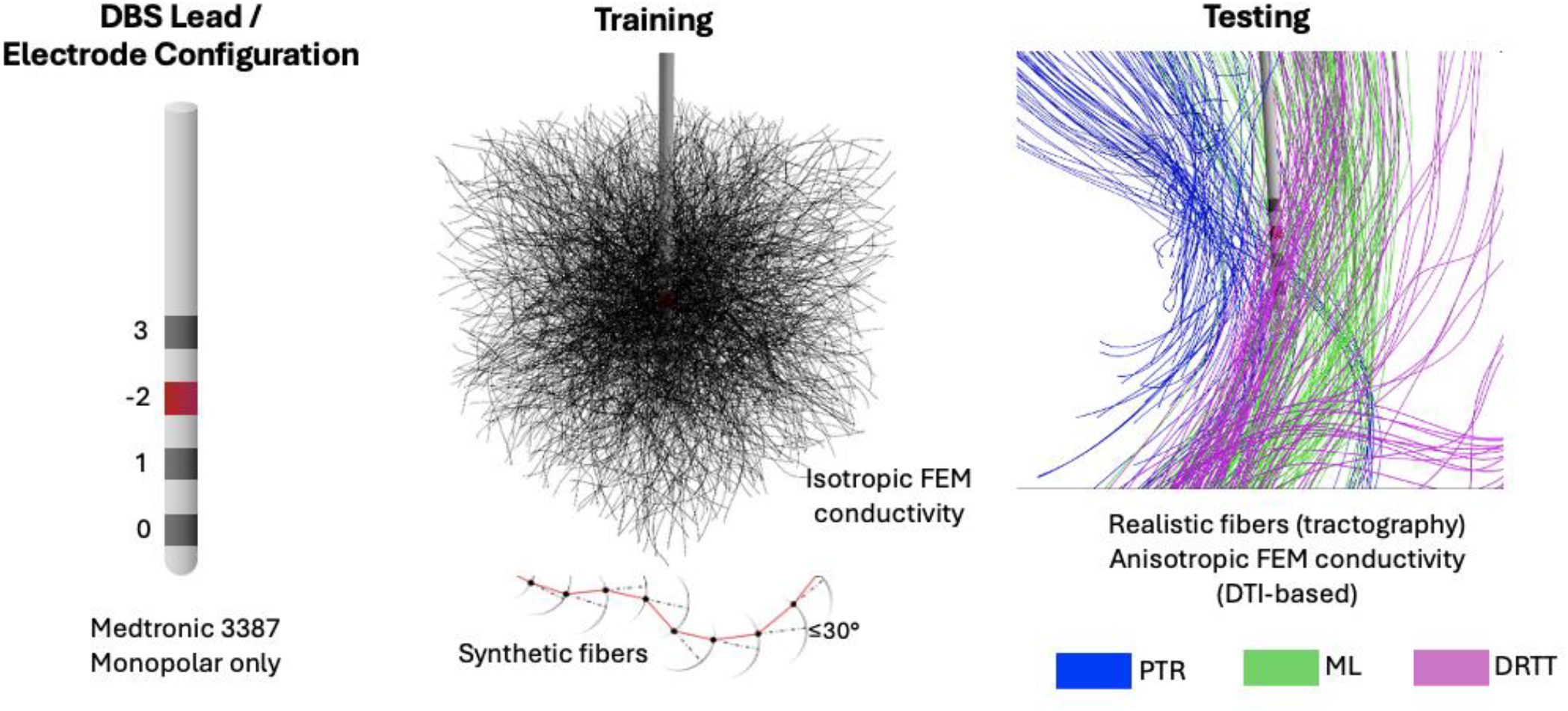
XGBoost training and testing FEM and fiber pathway conditions.

### 2.2 XGBoost Workflow

We aimed for a direct comparison between XGBoost and the best performing ANN model created by Golabek et al. [15]. To do this, we trained and tested XGBoost in the same way Golabek et al. trained and tested their best performing ANN model. Details are given in the Training and Testing sections below.

#### 2.2.1 XGBoost Training

We used NEURON/MRG-model to find the threshold voltage for 1000 synthetically generated fibers with 30º tortuosity, using a monopolar electrode in isotropic conductive media in the FEM (see Figure 1). Threshold voltages were generated across 9 pulse widths (60, 90, 120, 150, 200, 250, 300, 400, 500 µs). A classification and a regression training dataset was created from this data. For each fiber within the FEM bounds, a single row was made for the regression dataset and 100 rows were made for the classification dataset. To create these 100 variations for the classification dataset, each fiber’s spatial features were scaled linearly by 100 random stimulus amplitudes normally distributed around its true threshold. Variations meeting or exceeding the true threshold were labelled 1 (activated), and those below were labelled 0. Each row serves as a single training example comprising 35 variables: the stimulus pulse width, 33 spatial features (extracellular potentials, first spatial differences, and second spatial differences across an 11-node window), and a final target label (the exact activation threshold for regression, or a binary activation indicator for classification).

Hyperparameters are critical to the performance to a machine learning model and are not determined by training [32]. We determined the best hyperparameters by applying a grid search to 600 unique XGBoost hyperparameter combinations of interest. For each set of hyperparameters in the grid, a model was independently initialized and trained three times, and the average accuracy on the validation dataset (made from a random 20% of the training dataset) was used as the comparison metric. This was done separately on the classification and the regression training datasets. The final classification and regression XGBoost models were trained using their individual optimal hyperparameters. Each hyperparameter (number of boosting rounds, learning rate, max tree depth, subsample ratio, and feature subsampling) and their corresponding search scope can be found in Table 1.

**Table 1.**
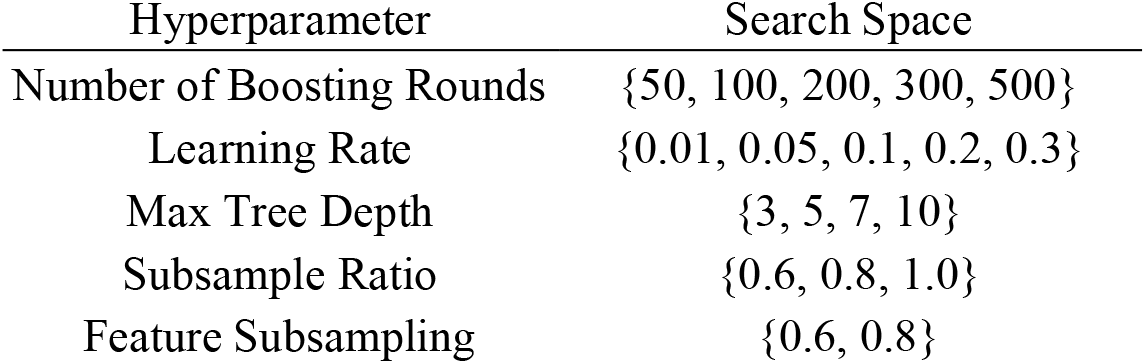
XGBoost Hyperparameter Optimization.

### 2.3 ANN and CNN Workflows

#### 2.3.1 Data Sets

We generated four datasets for both ANN And CNN models:

1. Training/validation for regression
2. Training/validation for classification
3. Testing for regression
4. Testing for classification

This separation was necessary because regression and classification tasks differ in their applicability to certain electrode configurations. In regression, activation thresholds must be scaled across all active contacts simultaneously. This scaling is meaningful for monopolar and bipolar cases, where only one contact (monopolar) or one voltage difference (bipolar) must be adjusted. In contrast, multipolar cases (e.g., double monopolar, triple monopolar, quadrupolar) cannot be meaningfully scaled for regression, since physicians can adjust individual contacts independently. Therefore, multipolar and asymmetric cases were restricted to classification experiments, where the task was to predict whether a given fiber was activated. The fiber tracts used for training/validation and testesting were the same as what was used for XGBoost training and testing as shown in Figure 1.

The Medtronic 3387 has 4 electrodes, ordered from distal to proximal and starting from 0. A +/-indicates that the corresponding electrode is active as an anode or cathode, respectively. If no electrode is configured as anodic, the implantable pulse generator (IPG) will be used as the anode, which will be designated as C+. For example, the double-monopolar configuration (0-3-C+) denotes that the first and fourth electrodes, counting from the tip of the lead up, are active (cathodic), while the second and third electrodes are inactive, and the IPG is used as the anode. Configurations for the BSC Vercise directional lead are labelled starting from one, and the middle two electrodes have three isolated segments (a,b,c). Plus and minus signs after any number/letter designates the anode and cathode electrodes, respectively. Figure 2 shows examples of the two DBS leads with different electrode configurations.

**Figure 2.**
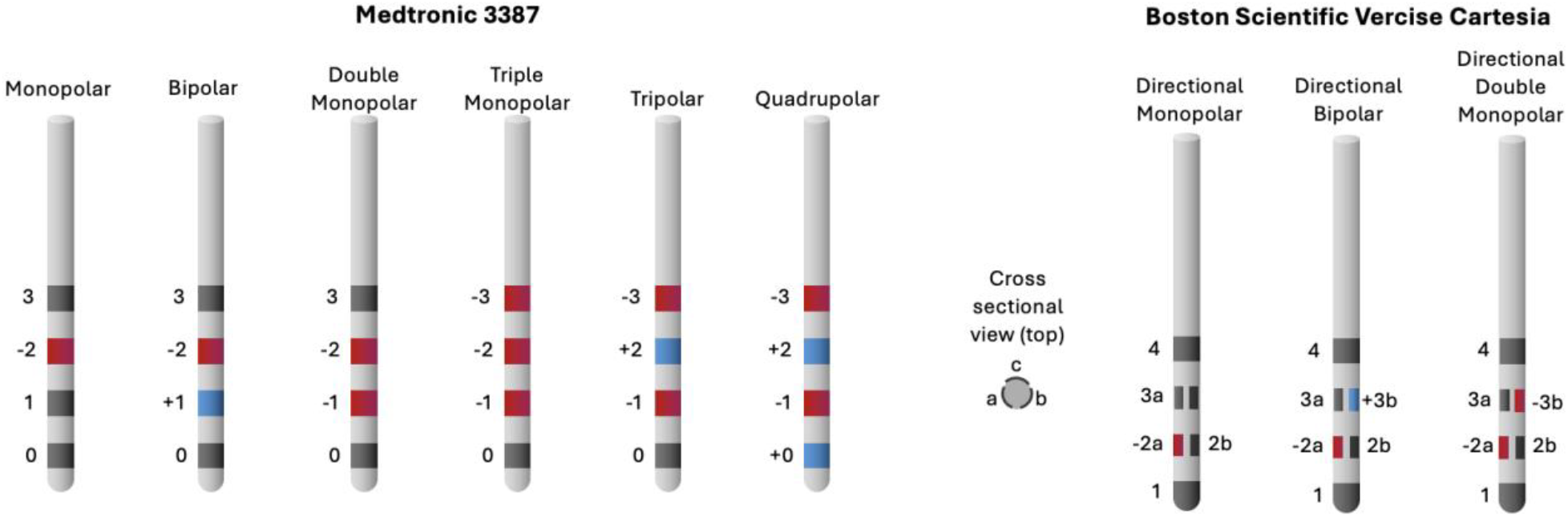
Medtronic and Boston Scientific DBS leads with example electrode configurations.

Regression models were trained on one Medtronic 3387 monopolar case (2-C+) and one Medtronic 3387 bipolar case (1-2+). Testing expanded beyond the two electrode configurations cases present in the training data, and also included monopolar and bipolar electrode cases for the BSC Vercise Cartesia directional lead (Table 2). Classification models were trained on a variety of electrode configurations for the Medtronic 3387 (Table 3). Testing included all regression test cases, plus multipolar and directional cases from both Medtronic and BSC systems, as well as several asymmetric cases on the Medtronic system (Table 4).

**Table 2.** Electrode testing set for regression ANNs and CNNs.

| Lead | Electrode Configuration | Case/Pattern |
| --- | --- | --- |
| Medtronic 3387 | Monopolar | 0- C+ |
| Medtronic 3387 | Monopolar | 1- C+ |
| Medtronic 3387 | Monopolar | 2- C+ |
| Medtronic 3387 | Monopolar | 3- C+ |
| Medtronic 3387 | Bipolar | 0- 3+ |
| Medtronic 3387 | Bipolar | 1- 2+ |
| Medtronic 3387 | Bipolar | 1+ 3- |
| BSC Vercise Cartesia | Directed Monopolar | 3a- C+ |
| BSC Vercise Cartesia | Directed Monopolar | 3b- C+ |
| BSC Vercise Cartesia | Directed Monopolar | 3c- C+ |
| BSC Vercise Cartesia | Directed Monopolar | 3ab- C+ |
| BSC Vercise Cartesia | Directed Monopolar | 3abc- C+ |
| BSC Vercise Cartesia | Directed Bipolar | 2a- 3a+ |
| BSC Vercise Cartesia | Directed Bipolar | 2b- 3a+ |
| BSC Vercise Cartesia | Directed Bipolar | 2ab- 3ab+ |
| BSC Vercise Cartesia | Directed Bipolar | 2ac- 3ab+ |
| BSC Vercise Cartesia | Directed Bipolar | 2abc- 3abc+ |

**Table 3.**
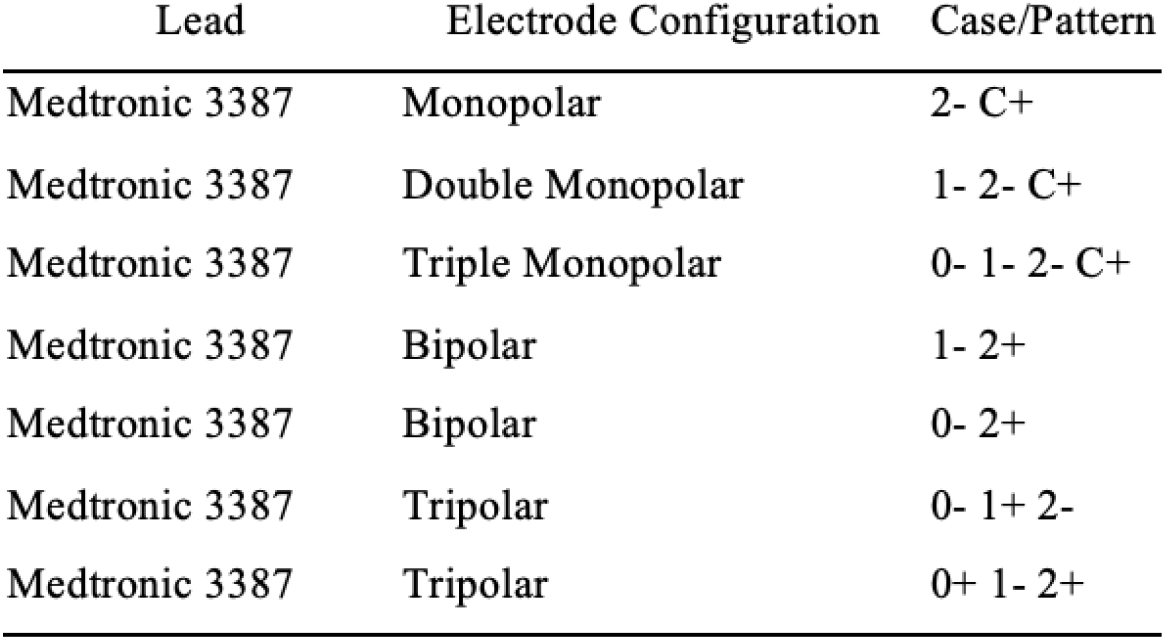
Electrode training/validation set for classification ANNs and CNNs.

| Lead | Electrode Configuration | Case/Pattern |
| --- | --- | --- |
| Medtronic 3387 | Monopolar | 2- C+ |
| Medtronic 3387 | Double Monopolar | 1- 2- C+ |
| Medtronic 3387 | Triple Monopolar | 0- 1- 2- C+ |
| Medtronic 3387 | Bipolar | 1- 2+ |
| Medtronic 3387 | Bipolar | 0- 2+ |
| Medtronic 3387 | Tripolar | 0- 1+ 2- |
| Medtronic 3387 | Tripolar | 0+ 1- 2+ |

**Table 4.** Electrode testing set exclusive to classification ANNs and CNNs. Electrode cases for regression (table 2) are also part of the classification test set.

| Lead | Electrode Configuration | Case/Pattern |
| --- | --- | --- |
| Medtronic 3387 | Double Monopolar | 1- 2- C+ |
| Medtronic 3387 | Double Monopolar | 0- 2- C+ |
| Medtronic 3387 | Double Monopolar | 0- 3- C+ |
| Medtronic 3387 | Triple Monopolar | 0- 1- 2- C+ |
| Medtronic 3387 | Triple Monopolar | 0- 1- 3- C+ |
| Medtronic 3387 | Quadruple Monopolar | 0- 1- 2- 3- C+ |
| Medtronic 3387 | Tripolar | 0- 1+ 2- |
| Medtronic 3387 | Tripolar | 0+ 1- 2+ |
| Medtronic 3387 | Tripolar | 0- 2- 3+ |
| Medtronic 3387 | Quadrupolar | 0- 1+ 2- 3+ |
| Medtronic 3387 | Quadrupolar | 0- 1+ 2+ 3- |
| Medtronic 3387 | Quadrupolar | 0+ 1- 2- 3+ |
| Medtronic 3387 | Asymmetric Bipolar | 0- 3+ |
| Medtronic 3387 | Asymmetric Bipolar | 1- 3+ |
| Medtronic 3387 | Asymmetric Double Monopolar | 0- 3- C+ |
| Medtronic 3387 | Asymmetric Double Monopolar | 1- 2- C+ |
| BSC Vercise Cartesia | Directed Double Monopolar | 2a- 3a- C+ |
| BSC Vercise Cartesia | Directed Double Monopolar | 2b- 3a- C+ |
| BSC Vercise Cartesia | Directed Double Monopolar | 2ab- 3ab- C+ |
| BSC Vercise Cartesia | Directed Double Monopolar | 2ac- 3ab- C+ |
| BSC Vercise Cartesia | Directed Double Monopolar | 2abc- 3abc- C+ |

Training and validation datasets for both regression and classification models were generated using synthetic fiber tracts from the random walk algorithm of [15], with a fiber tortuosity setting of 30° just like the XGBoost workflow (2.2). Test datasets were generated using DRTT, ML, and PTR fibers.

#### 2.3.2 Adjustments for Sites of Action Potential Initation

Previous DBS surrogate models [15] assumed that the node with the minimum extracellular voltage coincided with the site of action-potential initiation, and drew on previous findings that nodes further than 5 nodes away from the node of Ranvier made minimal contributions to the site of initiations [33]. Accordingly, an 11-node window centered on node with the minimum extracellular voltagewas used as input. For complex field geometries or tortuous fibers, this initiation site assumption may not hold. Similar to Wang et al. [16], we drew on Peterson et al.’s [34] use of weighted second spatial derivatives of the extracellular potential, where the node closest to the stimulation source has the heighest weight. To assess possible discrepancy between predicted site of initiation and MRG predicted site of initiation, the maximum second spatial difference (SSD) identified node was compared to the initiation site predicted by the MRG axon model for each fiber in our training dataset. From the resulting distributions (given in Supplemental Material), we determined the needed amount of nodes of Ranvier to cover 90%, 95%, and 99% of cases and converted it to a total window size, *W*. Larger windows capture a greater percentage of true initiation sites (as determined by MRG), but at the cost of increased input dimensionality.

Thus, to fully characterize the effect of spatial window size on ANN performance, we evaluated five input configurations for both regression and classification datasets:

Regression

1. Baseline: *W*=11nodes [15]
2. 90% coverage: *W* = 17 nodes
3. 95% coverage: *W* = 19 nodes
4. 99% coverage: *W* = 31 nodes
5. Large: *W* = 103 nodes

Classification

1. Baseline: *W*=11nodes [15]
2. 90% coverage: *W* = 17 nodes
3. 95% coverage: *W* = 21 nodes
4. 99% coverage: *W* = 39 nodes
5. Large: *W* = 103 nodes

In addition to the baseline and coverage-derived window sizes, we also evaluated a “Large” window of W = 103 nodes. This value was chosen to extend beyond the 99% coverage thresholds (31 and 39 nodes for regression and classification respectively) while still retaining ∼90% of the fibers available in the baseline W = 11 dataset. It serves as an upper-bound test of whether ANNs and CNNs could effectively handle substantially more spatial context or whether performance would plateau or degrade.

#### 2.3.3 ANN Model Parameters

ANN modeling followed the general workflow described by Golabek et al. [15], extended to incorporate multiple window sizes as defined in Section 2.3.2. This meant that networks were implemented as fully connected feedforward architectures, with input dimensionality set by the window size W. Inputs consisted of three features each of size W, i.e., extracellular potential, first spatial difference, and second spatial difference, plus the stimulus pulse width. Regression models were trained using mean absolute percentage error loss, and classification models using binary cross-entropy.

Hyperparameter optimization followed the same grid search technique that was present in [15], but restricted to ReLU activations. This decision was motivated by consistently poor results with tanh and sigmoid activations in preliminary experiments and by the well documented underperformance of these functions in multi-layer neural networks [35].

#### 2.3.4 CNN Model Parameters

CNN experiments were designed to test whether convolutional architectures could outperform ANNs by automatically learning spatial filters over the nodal profile. Architectures consisted of 1-D convolutional layers, each with a 50% probability of being followed by a max pooling layer, and were terminated with fully connected layers and shown in Figure 3.

**Figure 3.**
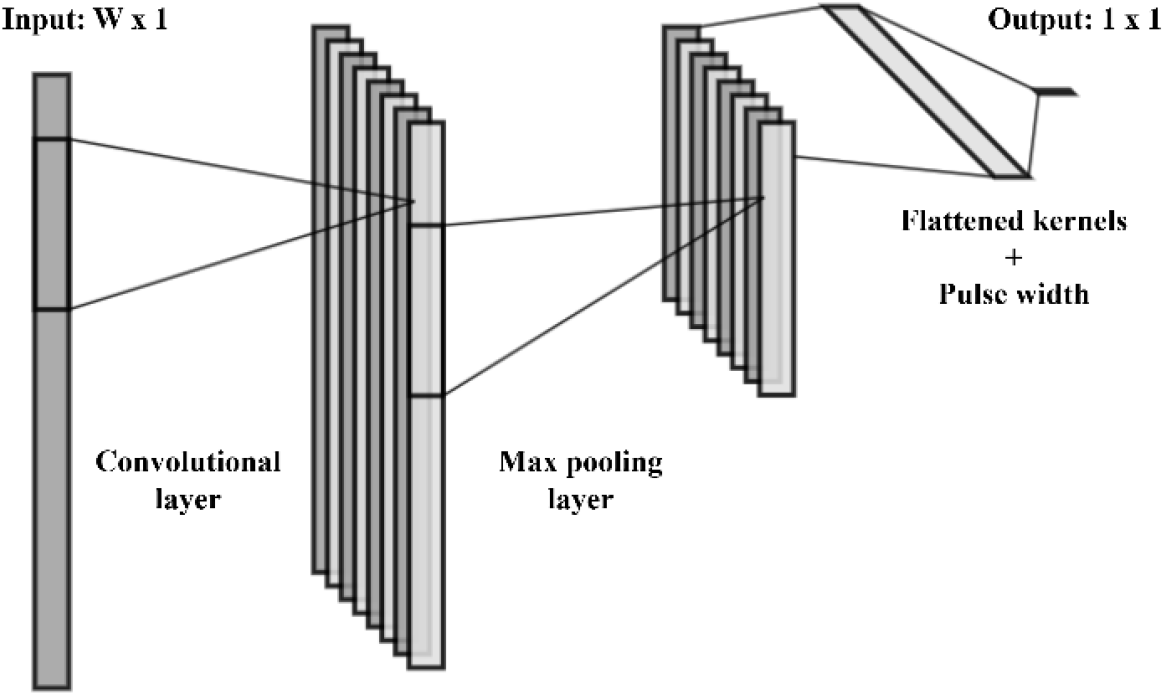
Example CNN Architecture. The example model includes all major layer types: input, convolution, max pooling, fully connected (dense), and output. For regression tasks the output is a positive scalar, and for classification tasks it is binary. Input length W is reduced after convolution due to TensorFlow’s default ‘valid’ padding, and the max pooling layer halves the dimensionality with default pool size of 2.

Hyperparameter optimization again followed the same grid search technique that was present in [15], while restricting to ReLU activations for the reasons described in Section 2.3.3. Now with CNN specific hyperparameters also incorporated. These CNN specific hyperparameters included the number of convolutional layers ranging from 1-4, kernel sizes of 3,5,7,9, or 11, and filter depths of 8, 16, 32, 64, or 128. All other parameters were set to TensorFlow 2.7 defaults.

Unlike ANNs, which were trained and evaluated on all of the windows specifications from 2.3, CNNs were trained and evaluated only on the two largest input windows (W=31/39 and W=103). This was done on the assumption that with an appropriate receptive field CNNs could essentially ‘block out the noise’ from the non-essential nodes. CNNs were also only given the extracellular potential, as we expect that the CNN would implicitly extract the SSD since it is a spatial operation.

#### 2.3.5 Training Procedure

All ANN and CNN models were trained for up to 100 epochs during hyperparameter searches with early stopping (patience = 10). The best performing hyperparameter configurations were then selected to train the final model based off the lowest validation set MAPE. In this final training process classification models were trained in the same manner, but regression models were allowed to run to a patience of 100 due to increasing progress noted in the hyperparameter search phase. Note that the hyperparameter search phase did not employ this larger training duration due to compute limitations from not being able to access GPUs. Models were trained and tested on AMD EPYC 7702 64-Core processors and AMD EPYC 75F3 32-Core processors.

#### 2.3.6 Testing procedure for XGBoost, ANN, and CNN models

XGBoost, ANN, and CNN models were tested on three white matter pathways derived from group-averaged DTI-based tractographies from the Human Connectome Project [31]. The three white matter pathways were from the posterior thalamic radiation (PTR), medial lemniscus (ML), and dentatorubrothalamic tract (DRTT) pathways (see Figure 1). The fibers were stimulated by an electrode in anisotropic, DTI-based conductivity conditions. The XGBoost model was only tested on a single monopolar electrode configuration, allowing for a direct comparison with Golabek et al. [15], while the ANN and CNN models were tested on many different electrode configurations. Testing was done on a set of pulse widths that included the pulse widths used in training, as well as a set of unseen pulse widths that were 50% between the training values (75, 105, 135, 175, 225, 275, 350, 450 µs) for 17 pulse widths tested in total.

To evaluate performance, the predicted threshold stimulus amplitudes (in volts) from the XGBoost, ANN, and CNN regression models were directly compared to the ground-truth MRG model predictions. Accuracy was quantified using mean absolute error (MAE) (|*V*_*th,model*_ - *V*_*th,MRG*_|) and mean absolute percent-error (MAPE) (|(*V*_*th,model*_ - *V*_*th,MRG*_(/*V*_*th,MRG*_ * 100%|). For classification models, threshold values were derived using the same binary search algorithm employed in the MRG simulations. This methodology ensures that we can use the same error metrics across all model architectures.

## 3. Results

We evaluated the performance of XGBoost, ANN, and CNN surrogate models in predicting nerve fiber activation across a range of clinical scenarios. Results are presented in three stages. First, we benchmark XGBoost against existing baselines to justify the selection of neural network architectures for complex fiber geometries. Second, we provide a comprehensive summary of ANN and CNN performance across diverse electrode configurations. Finally, we compare the computational runtimes of all models.

### 3.1 Comparing performance of XGBoost to ANN

To evaluate whether tree-based machine learning methods could match the predictive accuracy of deep learning architectures in complex neural environments, we benchmarked a classification and a regression XGBoost model against the gold-standard MRG axon model using three distinct white matter pathways (PTR, DRTT, and ML) under anisotropic conductivity conditions and compared the results to the best performing ANN of Golabek et al. [15]. The two XGBoost models were tested on a set of 100 PTR fibers, 100 DRTT fibers, and 100 ML fibers, as was the best performing ANN of Golabek et al. [15]

As shown in table 5, both the regression and the classification XGBoost models were outperformed by the best performing ANN model produced by Golabek et al. [15] However, the classification XGBoost model was clearly more accurate, and as such it will be the focus of the comparison of XGBoost to the ANN.

**Table 5.** XGBoost Results.

| XGBoost Classification | XGBoost Regression | Best performing ANN [15] |
| --- | --- | --- |
| MAPE (%) | MAPE (%) | MAPE (%) |
| 6.07 | 27.51 | 0.72 |

**Table 6.** Runtime comparison of various models on 288 fibers using the monopolar 2-C+ electrode.

| Model | Total Runtime (s) |
| --- | --- |
| Regression ANNs | 0.102 - 0.197 |
| Classification ANNs | 0.699 - 0.849 |
| Regression CNNs | 0.163 - 0.234 |
| Classification CNNs | 0.742 - 0.990 |
| Regression XGBoost | 0.024 |
| Classification XGBoost | 0.348 |
| MRG | 4638.776 |

Part of the reason why the XGBoost classification model performed poorly relative to the ANN was that it struggled to generalize to the 8 pulse widths that it was not trained on (Figure 4). Notably, the model exhibited a systematic overestimation of threshold voltages for unseen pulse widths, with the highest error magnitudes occurring at the shortest pulse durations.

**Figure 4.**
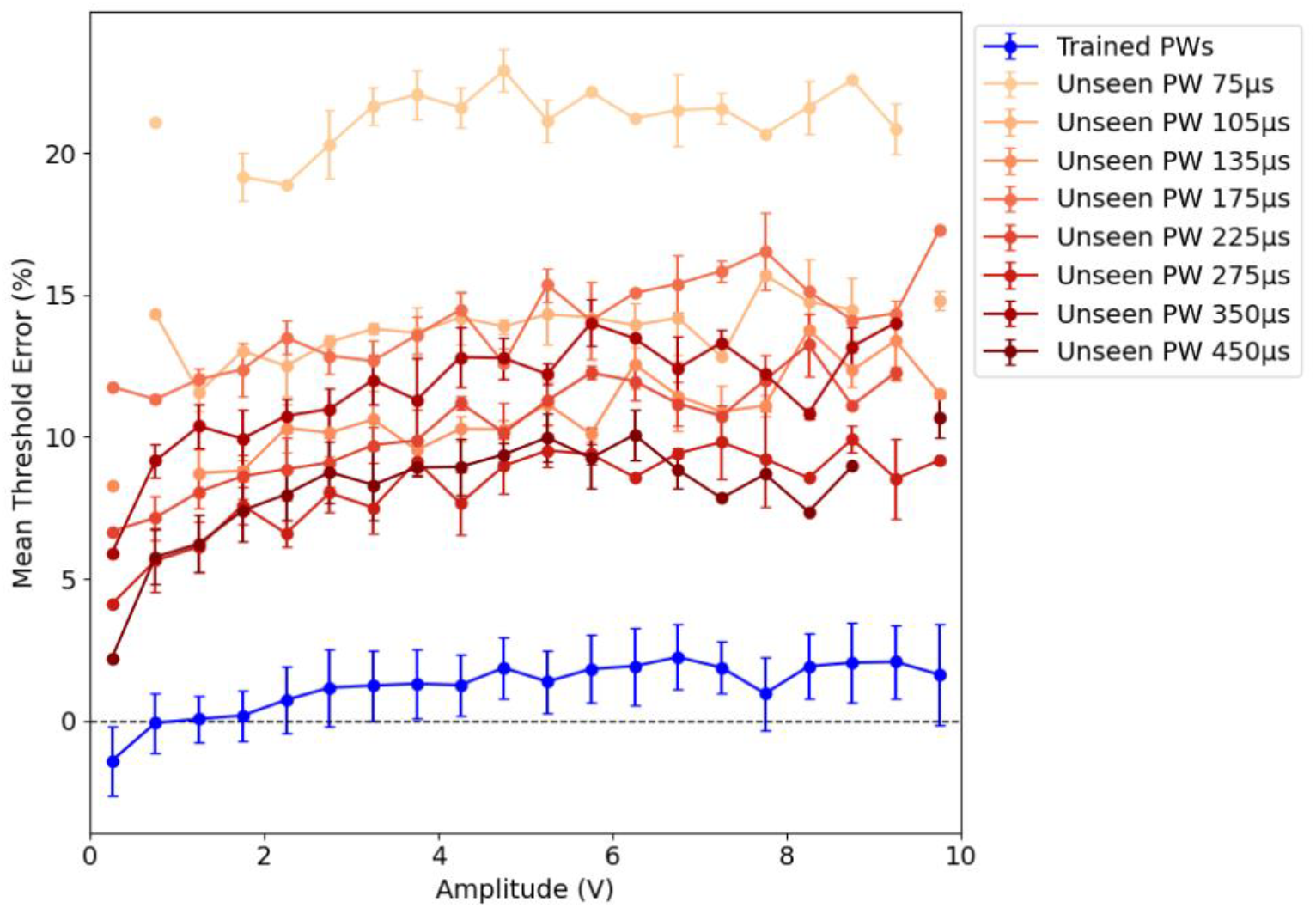
Percent error of the classification XGBoost model tested on PTR, DRTT, and ML fibers.

Given these limitations, we did not conduct the extended spatial window size analysis described in Section 2.3.2 for this architecture. Instead, we restricted XGBoost evaluation to this baseline configuration, as the preliminary results indicated it was not performant enough to warrant deeper window-size exploration.

### 3.2 Comparison of all ANN and CNN models

We evaluated the performance of ANN and CNN architectures across several spatial window configurations (ranging from W = 11 to W = 103 nodes). In clinical settings, the voltage applied by physicians is typically below 5 V in order to limit potential side effects [36,37]. As such, the ANN and CNN performance is evaluated in the 0 to 5 V range, where a comparison is made if either the model or MRG predicts a threshold voltage of less than 5 V. A corresponding current range to 0-5 V is 0-5.615 mA. Testing results are given in Figure 5.

**Figure 5.**
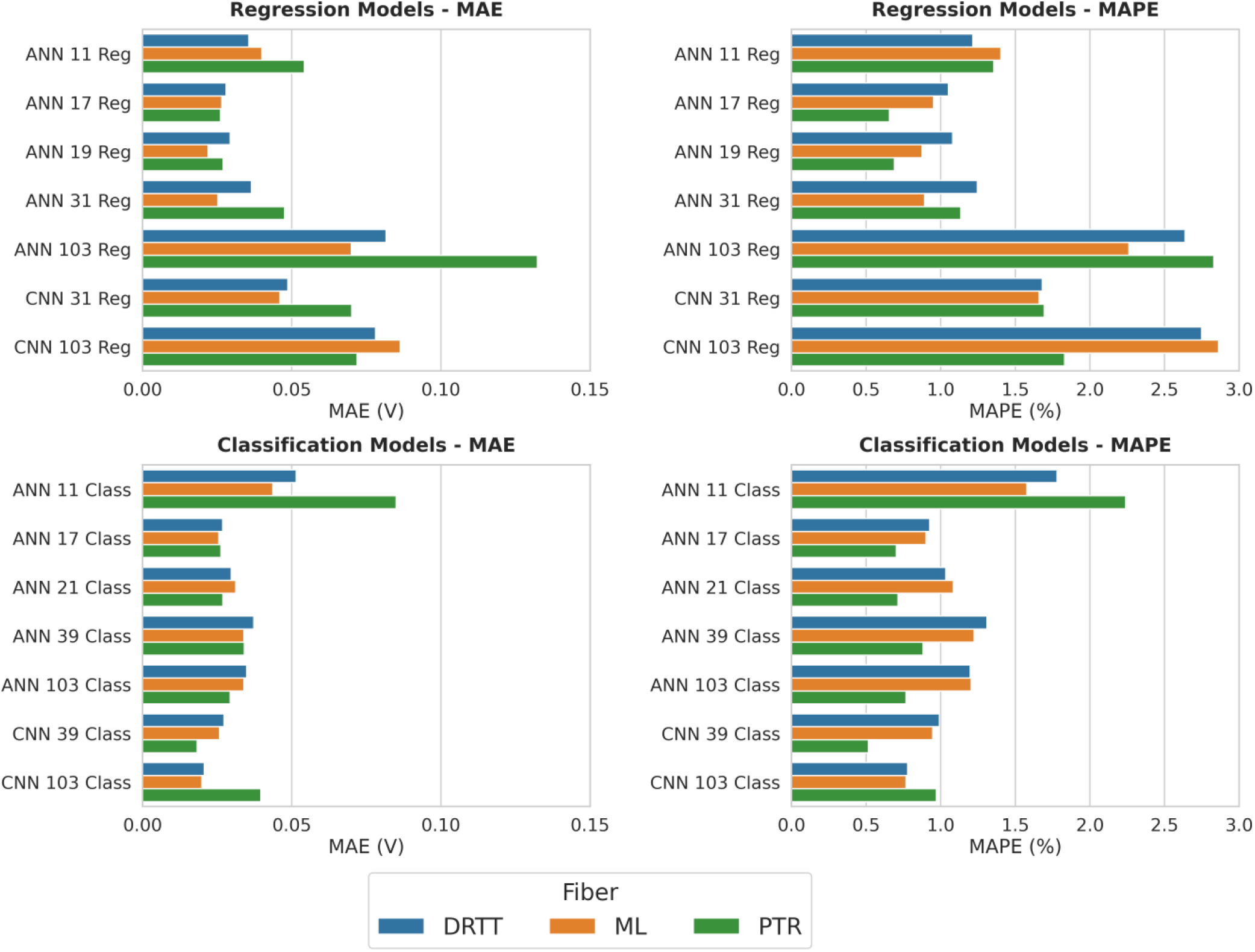
MAE and MAPE for all regression and classification models Across all regression models and fibers, the overall mean average error (MAE) was 0.0473 V, and the overall MAPE was 1.63%. Across all classification models and fibers, the overall MAE was 0.032 V and the overall MAPE was 1.13%. It should be noted that classification models are tested on more electrode configurations than regression models.

### 3.3 Comparison of all ANN and CNN Models by Electrode Cases

Analysis of individual electrode cases revealed that predictive accuracy is highly dependent on the specific electrode configuration of the DBS lead, the spatial windowing strategy employed by the model, and the architecture of the model. By evaluating the performance across the Medtronic 3387 and BSC families, we observe that different configurations favor specific architectures and window sizes. This section details the best performing models for each electrode configuration.

The classification CNN with a window size of 103 nodes was the best performing model on most configurations, with an overall MAPE and MAE of 0.77% and 0.0205 V, respectively (Figure 6 and Figure 7). The configurations that the classification CNN with a window size of 103 nodes did not perform the best on were directional bipolar, the quadruple monopolar subset of multipolar (although it was the best performing model overall for multipolar configurations), and asymmetric double monopolar, although it was still accurate on all three. The best performing models for these configurations were the classification ANN with a window size of 17 nodes, the classification CNN with a window size of 39 nodes, and the classification ANN with a window size of 21 nodes, respectively.

**Figure 6.**
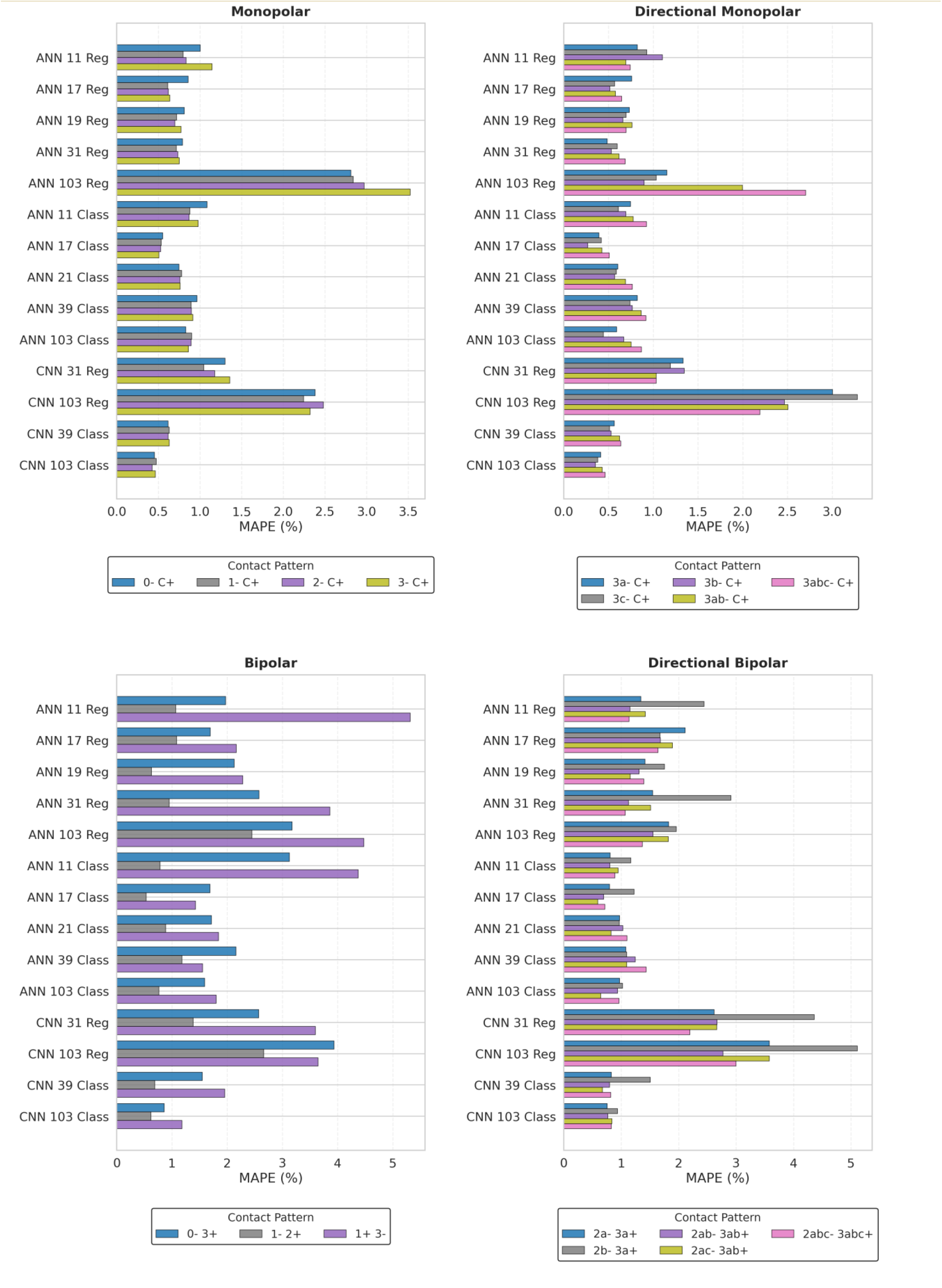
MAPE of classification and regression models across shared electrode configurations.

**Figure 7.**
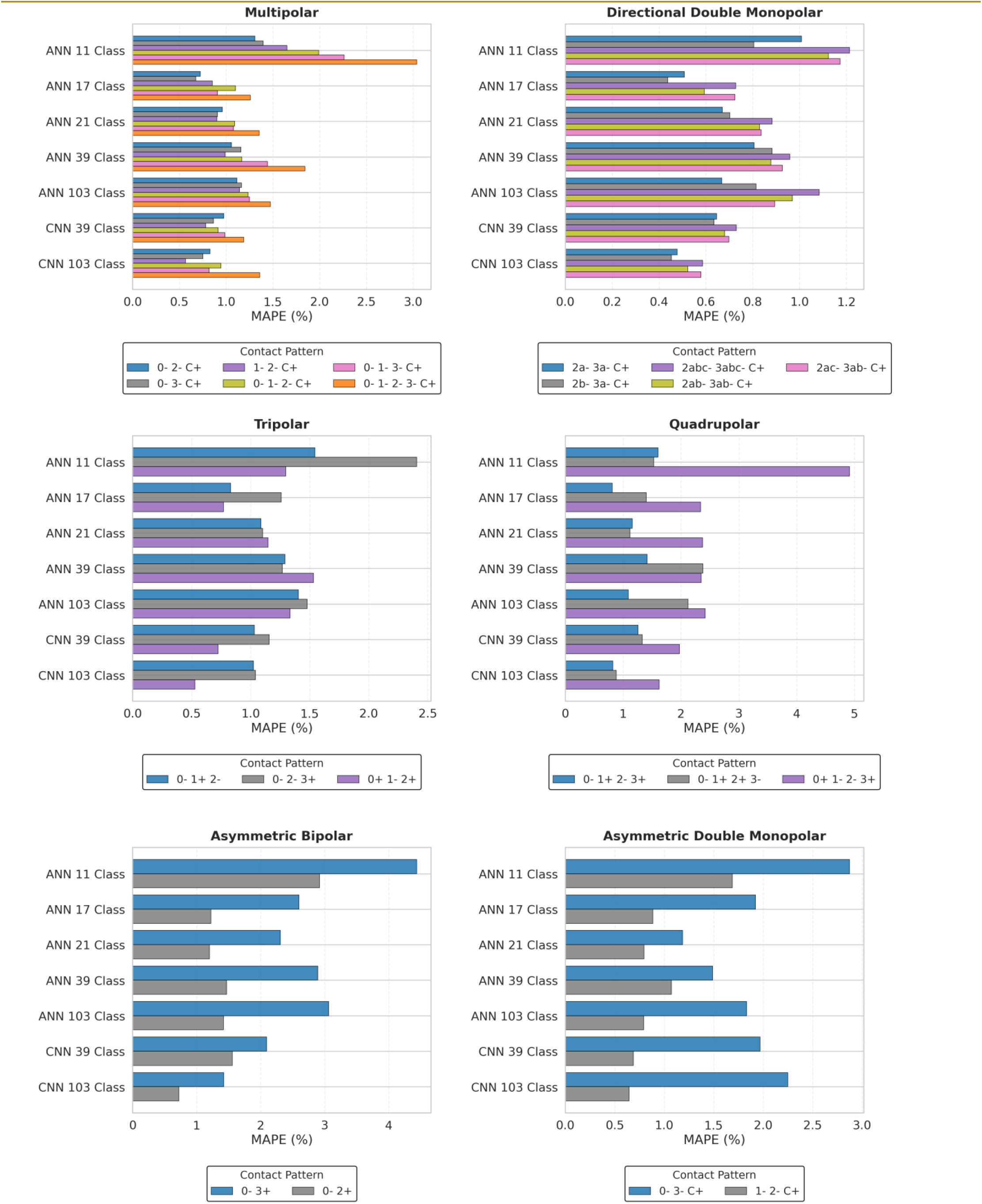
MAPE of classification models across electrode configurations not shared with regression models. Multipolar configurations include double monopolar, triple monopolar, and quadruple monopolar configurations. For asymmetric configurations, the more proximal (contact 3 is the most proximal electrode) contact is the one with the greater magnitude.

In terms of more general trends, both ANN and CNN classification models generally outperformed their regression counterparts. ANNs were the most accurate with a moderate expansion of window size, with the models that had window sizes of 17-21 nodes giving the best results for both classification and regression models. As for CNNs, the classification CNN with a window size of 103 nodes generally outperformed its 39 node counterpart, while the regression CNN with a window size of 31 nodes was more accurate than its counterpart with 103 nodes.

### 3.4 Runtime Comparison

The runtimes for all models using a Medtronic monopolar 2-C+ electrode configuration on 288 fibers are shown in Table 8. ANN, CNN, and XGBoost models decreased the time required to predict stimulus thresholds by roughly 4 to 5 orders of magnitude compared to the gold-standard MRG model. The regression XGBoost model was roughly four times faster than the fastest regression ANN, and the classification XGBoost model was roughly twice as fast as the fastest classification ANN. CNNs were only moderately slower than their ANN counterparts, and for ANNs and CNNs runtime typically increased as spatial window increased.

## 4. Discussion

### 4.1 XGBoost

#### 4.1.1 Classification Versus Regression

We found that the classification XGBoost model was significantly more accurate than the regression XGBoost model. Due to this large performance gap, only the classification model will be compared to the best performing ANN model produced by Golabek et al. [15].

#### 4.1.2 Comparison to ANN

We found that the classification XGBoost model was less accurate than the best performing ANN model created by Golabek et al. [15] with the classification XGBoost model having an MAPE of 6.07% and the ANN having an MAPE of 0.72%. This was partly because the classification XGBoost model was unable to generalize to pulse widths that it did not see in training, while Golabek et al. found no significant difference between pulse widths trained on and pulse widths only tested on. However, even when the unseen pulse widths were removed from the testing set, XGBoost still had a MAPE of 1.39%, roughly double the MAPE of the ANN. From this, we conclude that while boosted gradient trees such as XGBoost can perform well on straight fibers (i.e. peripheral nerves), as shown by Toni et al. [18], it is outperformed by ANN models when predicting the threshold voltage of complex tortuous fibers in anisotropic conductivity conditions of brain tissue.

#### 4.1.3 Generalizability

Figure 8 demonstrates that the classification XGBoost model almost uniformly overestimates threshold voltage. Furthermore, it showed that the magnitude of overestimation was significantly higher for pulse widths that XGBoost did not see in training compared to pulse widths that XGBoost had trained on, and that among the unseen pulse widths the magnitude of the error increased as the pulse width decreased. To explain this phenomenon, we theorize that the classification XGBoost model was unable to recognize that the strength duration curve of fiber activation is hyperbolic, and that XGBoost tried to generalize to unseen pulse widths linearly.

**Figure 8.**
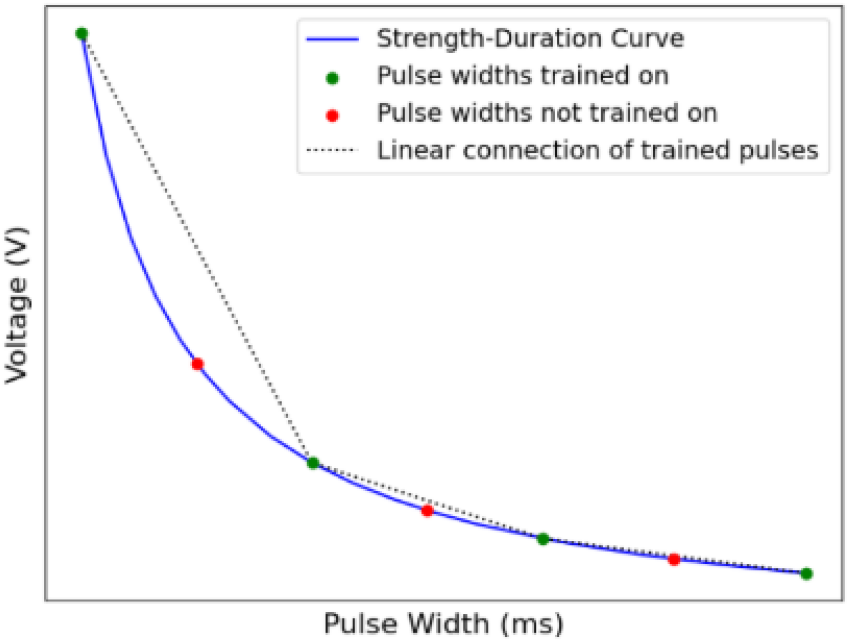
Strength-duration curve.

Figure 8 shows how generalizing linearly would lead to a consistent overestimation in threshold voltage. Due to the hyperbolic nature of the curve, when the pulse widths that are trained on are linearly connected the area under the linear connections is greater than the area under the original curve. This means that XGBoost would think that it takes more voltage than it actually does to activate fibers when using unseen pulse widths, and this effect would be worse for smaller unseen pulse widths because of the hyperbolic nature of the curve, which is consistent with our results. From this, we conclude that XGBoost cannot be relied upon to generalize to pulse widths not seen in training.

### 4.2 ANN and CNN models

ANNs and CNNs generalized successfully across monopolar, bipolar, tripolar, quadrupolar, multipolar, directional, and asymmetric configurations, demonstrating that surrogate models trained on a limited subset of electrode configurations can extend to more complex cases. This is a crucial fact given the combinatorial explosion that DBS programming faces.

Our analysis revealed several distinct patterns within the performance data. First, classification based ANNs and CNNs consistently outperformed their regression-based counterparts. This suggests that training models for the binary task of predicting activation at a specific voltage coupled with a binary search is a more effective strategy for future surrogate research than attempting to directly regress the threshold voltage. Secondly, we found that the classification CNN with a window size of 103 nodes was the best performing model overall. It was the most accurate model on all electrode configurations except for directional bipolar and asymmetric double monopolar, yet it remained robust on those configurations as well. These results corroborate findings by Wang et al. [16], suggesting that CNNs are the best equipped architecture for prediction of single fiber activation in DBS.

### 4.3 Study Limitations

While generalization across electrode families and configurations were tested, all training data was derived from FEMs under controlled conditions, limiting the ability to make any definitive claim on generalization. Moreover, prediction results are defined for 5.7 mm diameter fibers, only.

### 4.4 Future Directions

Future research could test other CNN architectures. Of particular interest would be architectures where an inductive bias was inserted into the model in the form of separate convolutional branches that come together during pulse-width concatenation, essentially functioning as untrainable kernels that the CNN could leverage for possible faster convergence or improved accuracy. Additionally, this study looked at ANNs and CNNs as the main deep learning methods, but attention-based methods are worth exploring, especially those which are more able to pick up on short term and bidirectional relationships between inputs. This would entail exploring model performance for bidirectional Long Short Term Memory (LSTM) networks and Gated Recurrent Units (GRUs). Finally, training data could be improved to extend it to different fiber diameters and other stimulus waveforms for additional modelling ability.

## 5. Conclusions

We developed and compared ANN- and CNN-based surrogate models for prediction of DBS single fiber activation that extend beyond monopolar stimulation, to include bipolar, tripolar, quadrupolar, multipolar, directional monopolar, directional bipolar, and directional double monopolar, as well as several asymmetric configurations. We evaluated the machine learning predictors using white matter pathways derived from group-averaged connectome data within a patient-specific tissue conductivity field, comparing both predicted stimulus activation thresholds and pathway recruitment across a clinically relevant range of stimulus amplitudes and pulse widths. We compared 14 different ANN or CNN models that performed either classification or regression and varied in the amount of extracellular voltage data input to the model. In particular, the classification CNN with a window size of 103 nodes (i.e., the voltage data input to the model comprised the predicted node of initiation plus 51 Nodes of Ranvier on either side) emerged as the overall most accurate and robust predictor of all the models tested.

We also compared XGBoost models to ANN models for monopolar electrode configurations, only. XGBoost in contrast to ANNs, had a relatively high error for a simpler task of modelling monopolar stimulation. These results highlight the limitations of classical machine learning methods in settings such as DBS, where untrained for heterogeneity may arise, and reinforce the advantages of deep learning methods such as ANN- and CNN-based approaches for DBS surrogate modelling. Together, this work advances the development of surrogate predictors for DBS by expanding to tripolar, quadrupolar, multipolar, and asymmetric configurations for classification, improving generalization across electrodes and electrode configurations, and enhancing interpretability. These contributions strengthen the case for adopting machine learning-based tools in clinical DBS programming.

## Data

All trained ANN and CNN models, along with the voltage field profiles for electrode configurations and fiber pathways used in this study, can be accessed and tested through this GitHub repository: https://github.com/Erinpatrick/DBS_SingleFiber_Rapid_Predictor_2.0.

## References

1. Lozano AM, Lipsman N, Bergman H, Brown P, Chabardes S, Chang JW, et al. Deep brain stimulation: current challenges and future directions. Nat Rev Neurol. 2019 Mar;15(3):148–60. doi:10.1038/s41582-018-0128-2 PubMed PMID: 30683913; PubMed Central PMCID: PMC6397644.

2. Martinez-Nunez AE, Rozell CJ, Little S, Tan H, Schmidt SL, Grill WM, et al. Proceedings of the 12th annual deep brain stimulation think tank: cutting edge technology meets novel applications. Front Hum Neurosci. 2025 Feb 25;19. doi:10.3389/fnhum.2025.1544994

3. Lozano AM, Lipsman N. Probing and Regulating Dysfunctional Circuits Using Deep Brain Stimulation. Neuron. 2013 Feb 6;77(3):406–24. doi:10.1016/j.neuron.2013.01.020 PubMed PMID: 23395370.

4. Vitek JL, Jain R, Chen L, Tröster AI, Schrock LE, House PA, et al. Subthalamic nucleus deep brain stimulation with a multiple independent constant current-controlled device in Parkinson’s disease (INTREPID): a multicentre, double-blind, randomised, sham-controlled study. The Lancet Neurology. 2020 Jun 1;19(6):491–501. doi:10.1016/S1474-4422(20)30108-3

5. Krauss JK, Lipsman N, Aziz T, Boutet A, Brown P, Chang JW, et al. Technology of deep brain stimulation: current status and future directions. Nat Rev Neurol. 2021 Feb;17(2):75–87. doi:10.1038/s41582-020-00426-z PubMed PMID: 33244188; PubMed Central PMCID: PMC7116699.

6. Wong JK, Middlebrooks EH, Grewal SS, Almeida L, Hess CW, Okun MS. A Comprehensive Review of Brain Connectomics and Imaging to Improve Deep Brain Stimulation Outcomes. Movement Disorders. 2020;35(5):741–51. doi:10.1002/mds.28045

7. Butson CR, Cooper SE, Henderson JM, Wolgamuth B, McIntyre CC. Probabilistic analysis of activation volumes generated during deep brain stimulation. NeuroImage. 2011 Feb 1;54(3):2096–104. doi:10.1016/j.neuroimage.2010.10.059

8. Cheung T, Noecker AM, Alterman RL, McIntyre CC, Tagliati M. Defining a therapeutic target for pallidal deep brain stimulation for dystonia. Annals of Neurology. 2014 Jul 1;76(1):22–30. doi:10.1002/ana.24187

9. Eisenstein SA, Koller JM, Black KD, Campbell MC, Lugar HM, Ushe M, et al. Functional anatomy of subthalamic nucleus stimulation in Parkinson disease. Annals of Neurology. 2014;76(2):279–95. doi:10.1002/ana.24204

10. Dembek TA, Barbe MT, Åström M, Hoevels M, Visser-Vandewalle V, Fink GR, et al. Probabilistic mapping of deep brain stimulation effects in essential tremor. NeuroImage: Clinical. 2017 Jan 1;13:164–73. doi:10.1016/j.nicl.2016.11.019

11. Reich MM, Horn A, Lange F, Roothans J, Paschen S, Runge J, et al. Probabilistic mapping of the antidystonic effect of pallidal neurostimulation: a multicentre imaging study. Brain. 2019 May 1;142(5):1386–98. doi:10.1093/brain/awz046

12. Horn A, Ostwald D, Reisert M, Blankenburg F. The structural-functional connectome and the default mode network of the human brain. Neuroimage. 2014 Nov 15;102 Pt 1:142–51. doi:10.1016/j.neuroimage.2013.09.069 PubMed PMID: 24099851.

13. Al-Fatly B, Ewert S, Kübler D, Kroneberg D, Horn A, Kühn AA. Connectivity profile of thalamic deep brain stimulation to effectively treat essential tremor. Brain. 2019 Oct 1;142(10):3086–98. doi:10.1093/brain/awz236

14. Patrick EE, Fleeting CR, Patel DR, Casauay JT, Patel A, Shepherd H, et al. Modeling the volume of tissue activated in deep brain stimulation and its clinical influence: a review. Frontiers in Human Neuroscience [Internet]. 2024;18. Available from: https://www.frontiersin.org/articles/10.3389/fnhum.2024.1333183

15. Golabek J, Schiefer M, Wong JK, Saxena S, Patrick E. Artificial neural network-based rapid predictor of biological nerve fiber activation for DBS applications. Journal of Neural Engineering. 2023 Jan 18;20(1):016001. doi:10.1088/1741-2552/acb016

16. Wang S, Ma R, Yuan Q, Li H, Jiang C. Efficient, Robust, and Accurate CNN Predictor for Neuronal Activation in Directional Deep Brain Stimulation. IEEE Trans Neural Syst Rehabil Eng. 2025;33:1685–94. doi:10.1109/TNSRE.2025.3561122 PubMed PMID: 40232895.

17. Li H, Wang S, Luo X, Jiang C, Zhang B. Versatile Neural Activation Predictor With Axon Structure Tailoring Capability Enabling Personalized Neuromodulation Computation. IEEE Transactions on Neural Systems and Rehabilitation Engineering. 2025;33:3986–97. doi:10.1109/TNSRE.2025.3614215

18. Toni L, Pierantoni L, Verardo C, Romeni S, Micera S. Characterization of Machine Learning-Based Surrogate Models of Neural Activation Under Electrical Stimulation. Bioelectromagnetics. 2025 Jan;46(1):e22535. doi:10.1002/bem.22535

19. Grill WM, Mortimer JT. Electrical properties of implant encapsulation tissue. Ann Biomed Eng. 1994;22(1):23–33. doi:10.1007/BF02368219 PubMed PMID: 8060024.

20. Gunalan K, Howell B, McIntyre CC. Quantifying axonal responses in patient-specific models of subthalamic deep brain stimulation. NeuroImage. 2018 May 15;172:263–77. doi:10.1016/j.neuroimage.2018.01.015

21. Howell B, Gunalan K, McIntyre CC. A Driving-Force Predictor for Estimating Pathway Activation in Patient-Specific Models of Deep Brain Stimulation. Neuromodulation: Technology at the Neural Interface. 2019 Jun 1;22(4):403–15. doi:10.1111/ner.12929

22. Haueisen J, Tuch DS, Ramon C, Schimpf PH, Wedeen VJ, George JS, et al. The Influence of Brain Tissue Anisotropy on Human EEG and MEG. NeuroImage. 2002 Jan 1;15(1):159–66. doi:10.1006/nimg.2001.0962

23. Plonsey R, Heppner DB. Considerations of quasi-stationarity in electrophysiological systems. Bulletin of Mathematical Biophysics. 1967 Dec 1;29(4):657–64. doi:10.1007/BF02476917

24. McIntyre CC, Richardson AG, Grill WM. Modeling the excitability of mammalian nerve fibers: influence of afterpotentials on the recovery cycle. J Neurophysiol. 2002 Feb;87(2):995–1006. doi:10.1152/jn.00353.2001 PubMed PMID: 11826063.

25. Hines ML, Carnevale NT. NEURON: a tool for neuroscientists. Neuroscientist. 2001 Apr;7(2):123–35. doi:10.1177/107385840100700207 PubMed PMID: 11496923.

26. Moffitt MA, Mcintyre CC, Grill WM. Prediction of myelinated nerve fiber stimulation thresholds: limitations of linear models. IEEE Transactions on Biomedical Engineering. 2004 Feb;51(2):229–36. doi:10.1109/TBME.2003.820382

27. Freeberg MJ, Schiefer MA, Triolo RJ. Efficient search and fit methods to find nerve stimulation parameters for multi-contact electrodes. In: 2011 Annual International Conference of the IEEE Engineering in Medicine and Biology Society [Internet]. 2011 [cited 2026 Jan 24]. p. 7238–41. Available from: https://ieeexplore.ieee.org/abstract/document/6091829 doi:10.1109/IEMBS.2011.6091829

28. Yeh FC, Panesar S, Fernandes D, Meola A, Yoshino M, Fernandez-Miranda JC, et al. Population-averaged atlas of the macroscale human structural connectome and its network topology. NeuroImage. 2018 Sep 1;178:57–68. doi:10.1016/j.neuroimage.2018.05.027

29. Setsompop K, Kimmlingen R, Eberlein E, Witzel T, Cohen-Adad J, McNab JA, et al. Pushing the limits of in vivo diffusion MRI for the Human Connectome Project. NeuroImage. 2013 Oct 15;Mapping the Connectome 80:220–33. doi:10.1016/j.neuroimage.2013.05.078

30. Fan Q, Witzel T, Nummenmaa A, Van Dijk KRA, Van Horn JD, Drews MK, et al. MGH–USC Human Connectome Project datasets with ultra-high b-value diffusion MRI. NeuroImage. 2016 Jan 1;Sharing the wealth: Brain Imaging Repositories in 2015124:1108–14. doi:10.1016/j.neuroimage.2015.08.075

31. Van Essen DC, Ugurbil K, Auerbach E, Barch D, Behrens TEJ, Bucholz R, et al. The Human Connectome Project: a data acquisition perspective. Neuroimage. 2012 Oct 1;62(4):2222–31. doi:10.1016/j.neuroimage.2012.02.018 PubMed PMID: 22366334; PubMed Central PMCID: PMC3606888.

32. Probst P, Boulesteix AL, Bischl B. Tunability: Importance of Hyperparameters of Machine Learning Algorithms. Journal of Machine Learning Research. 2019;20(53):1–32.

33. Warman EN, Grill WM, Durand D. Modeling the effects of electric fields on nerve fibers: determination of excitation thresholds. IEEE Trans Biomed Eng. 1992 Dec;39(12):1244–54. doi:10.1109/10.184700 PubMed PMID: 1487287.

34. Peterson EJ, Izad O, Tyler DJ. Predicting myelinated axon activation using spatial characteristics of the extracellular field. J Neural Eng. 2011 Jul;8(4):046030. doi:10.1088/1741-2560/8/4/046030

35. Glorot X, Bordes A, Bengio Y. Deep Sparse Rectifier Neural Networks. In: Proceedings of the Fourteenth International Conference on Artificial Intelligence and Statistics. JMLR Workshop and Conference Proceedings; 2011. p. 315–23.

36. Volkmann J. Deep Brain Stimulation for the Treatment of Parkinson’s Disease. Journal of Clinical Neurophysiology. 2004;21(1).

37. Neumann WJ, Steiner LA, Milosevic L. Neurophysiological mechanisms of deep brain stimulation across spatiotemporal resolutions. Brain. 2023 Nov 1;146(11):4456–68. doi:10.1093/brain/awad239

