## Supplemental Data for "Comparison of Machine Learning Surrogate Models for Prediction of Single-Fiber Activation in Deep Brain Stimulation"

### ***Investigating maximum second spatial difference as an accurate predictor of initiation site for ANNs and CNNs***

Analysis revealed that the 11-node input window did not always align with the true action potential initiation site from electrical stimulation. To capture 90% of the true initiation sites a 17-node input window was needed, for 95% and 99% this number was 21 and 39 respectively for the classification dataset, and 19 and 31 respectively for the regression dataset. The initiation site offset ($\Delta_{init}$) quantifies the difference between the node of maximum second spatial difference (SSD) and is either the initiation site node index predicted by MRG or the mean of all initiation site node indices predicted by the MRG axon model. When multiple initiation sites were present in the MRG output, their indices were averaged.

The results are illustrated below for the classification dataset in the form of histograms (Figure 1), detailing the error between the two methods for determining the initiation site, and boxplots in Figure 2, displaying the same information sectioned off by the specific electrode configuration and case.

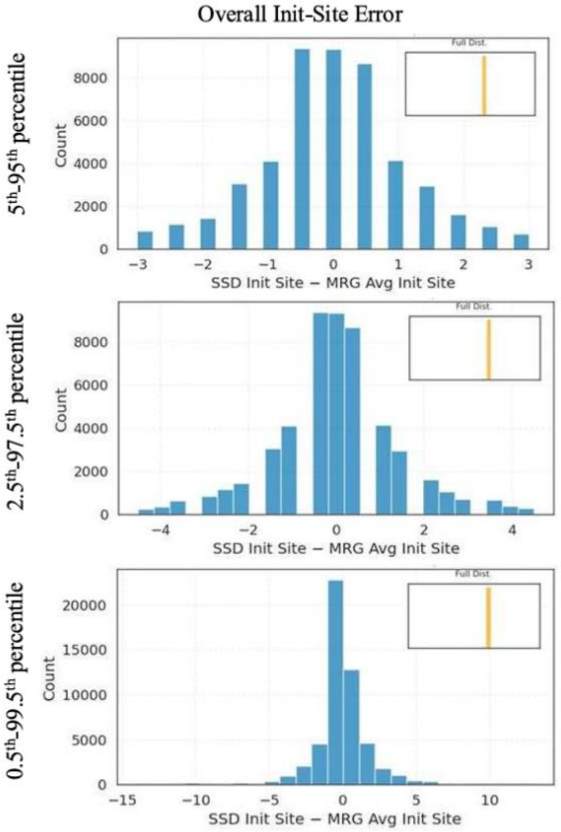


**Figure 1.** Distribution of the initiation-site offset (SSD Init Site – MRG Avg Init Site) shown for different percentile ranges

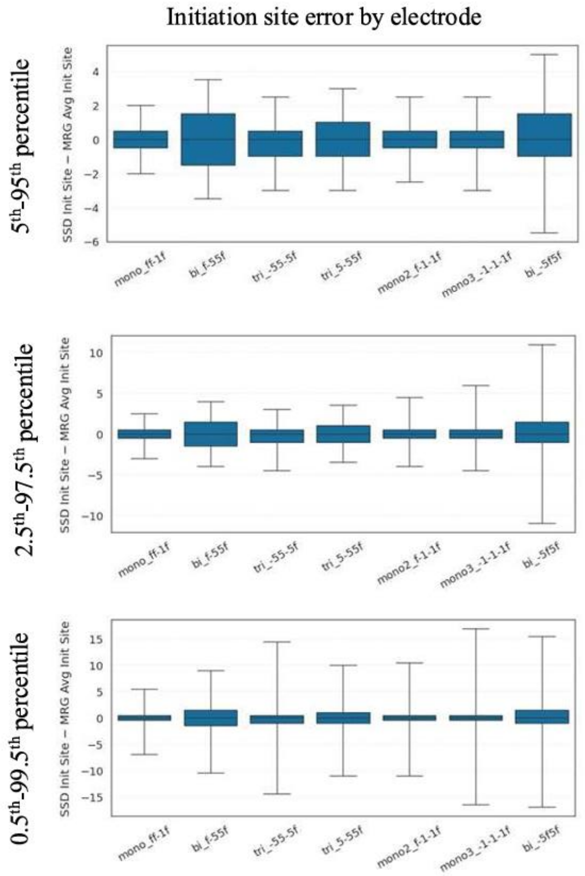


**Figure 2.** Distribution of the initiation-site offset (SSD Init Site – MRG Avg Init Site) shown for different percentile ranges by electrode configuration

### *Testing of ANN and CNN Models*

The tables below show the number of realistic fibers in each tract that were used to test the accuracy of the ANN and CNN models. Only fibers that were activated within the stimulus voltage window of 0-5V were included. As one can see, the number of fibers used in the PTR tract were significantly less than the other tracts.

***Table 6.*** *Number of comparisons by regression model and fiber tract.*

| Model | DRTT | ML | PTR | Total |
|---|---|---|---|---|
| ANN 11 | 4514 | 2220 | 58 | 6792 |

| | | | | |
|---|---|---|---|---|
| ANN 17 | 4506 | 2218 | 58 | 6782 |
| ANN 19 | 4525 | 2220 | 58 | 6803 |
| ANN 31 | 4530 | 2225 | 58 | 6813 |
| ANN 103 | 4480 | 2217 | 3 | 6700 |
| CNN 31 | 4515 | 2223 | 58 | 6796 |
| CNN 103 | 4534 | 2258 | 3 | 6795 |

***Table 7.*** *Number of comparisons by classification model and fiber tract.*

| Model | DRTT | ML | PTR | Total |
|---|---|---|---|---|
| ANN 11 | 12739 | 7297 | 984 | 21020 |
| ANN 17 | 12716 | 7291 | 975 | 20982 |
| ANN 21 | 12736 | 7316 | 955 | 21007 |
| ANN 39 | 12678 | 6949 | 640 | 20267 |
| ANN 103 | 12600 | 7241 | 132 | 19973 |
| CNN 39 | 12722 | 7284 | 976 | 20982 |
| CNN 103 | 12687 | 7277 | 977 | 20941 |
